# Effects of Emodin, a Plant-Derived Anthraquinone, on TGF-β1-Induced Cardiac Fibroblast Activation and Function

**DOI:** 10.1101/2021.01.07.425762

**Authors:** Wayne Carver, Ethan Fix, Charity Fix, Daping Fan, Mrinmay Chakrabarti, Mohamad Azhar

## Abstract

Cardiac fibrosis accompanies a number of pathological conditions and results in altered myocardial structure, biomechanical properties and function. The signaling networks leading to fibrosis are complex, contributing to the general lack of progress in identifying effective therapeutic approaches to prevent or reverse this condition. Several studies have shown protective effects of emodin, a plant-derived anthraquinone, in animal models of fibrosis. A number of questions remain regarding the mechanisms whereby emodin impacts fibrosis. TGF-β1 is a potent stimulus of fibrosis and fibroblast activation. In the present study, experiments were performed to evaluate the effects of emodin on activation and function of cardiac fibroblasts following treatment with TGF-β1. We demonstrate that emodin attenuates TGF-β1-induced fibroblast activation and collagen accumulation *in vitro*. Emodin also inhibits activation of several canonical (SMAD2/3) and non-canonical (Erk1/2) TGF-β signaling pathways, while activating the p38 pathway. These results suggest that emodin may provide an effective therapeutic agent for fibrosis that functions via specific TGF-β signaling pathways.

## Introduction

Fibrosis, characterized by excessive accumulation of extracellular matrix and matricellular proteins, is a hallmark of adverse myocardial remodeling and accompanies many pathological conditions of the heart (Spinale et al., 2016; Frangogiannis, 2019a). In the heart, fibrosis results in altered ventricular structure, geometry and biomechanical properties, which contribute to the progression of heart failure. Fibrosis in the heart can accompany massive loss of cardiomyocytes for instance following myocardial infarction in what has been termed “replacement fibrosis”. Myocardial fibrosis can also occur as a consequence of the direct activation of fibroblasts in the absence of substantial cardiomyocyte death by several stressors including pressure overload, diabetes, alcohol abuse and others. The progression of fibrosis often contributes to organ dysfunction and failure not only in the heart but in other organs as well. Relatively recent studies suggesting that fibrosis may be reversible, at least in its early stages, have stimulated interest in identifying potential therapeutic targets of fibrosis and activation of extracellular matrix-producing cells (Kumar and Sarin, 2007; Frangogiannis, 2019b).

The formation of specialized cells termed myofibroblasts is an important step in the progression of fibrosis. These cells are important mediators of normal wound healing but are also involved in pathogenesis of fibrosis and other conditions (Hinz, 2016; Shu and Lovicu, 2017; Pakshir and Hinz, 2018). As their name suggests, myofibroblasts have properties of smooth muscle cells and fibroblasts. They have enhanced synthetic activity producing extracellular matrix components, cytokines and growth factors and generate contractile force on their extracellular matrix (Carthy, 2018; Pakshir and Hinz, 2018). In the heart, these cells are potentially derived from multiple sources including: 1) resident fibroblasts that transform or differentiate in response to cardiac injury or stress, 2) epicardial or endothelial cells that undergo epithelial- or endothelial-to-mesenchymal transition, respectively or 3) potentially extra-cardiac stem cells (Kanisicak et al., 2016; Travers et al., 2016). Despite their origin, recent studies targeting these cells have conclusively identified their roles in myocardial fibrosis and have provided substantial insight into the molecular mechanisms of this process (Khalil et al., 2017; Bhandary et al., 2018).

TGF-β and its signaling pathways are important contributors to fibrosis generated by a number of stressors (Branton and Hoop, 1999; Biernacka et al., 2011; Leask and Abraham, 2004; Khalil et al., 2017). TGF-β is a multifunctional growth factor that modulates diverse cellular processes including proliferation, gene expression, migration, cell death and others (Derynck and Budi, 2019). Due to its pleiotropic cellular effects, it is not surprising that dysregulation of TGF-β signaling is involved in a number of disease processes including not only fibrosis, but cancer, autoimmunity and others. The canonical TGF-β signaling cascade is initiated by ligand binding to its receptor complex and recruitment of receptor-activated SMAD proteins (SMAD2 and SMAD3). SMAD2 and SMAD3 are activated by the receptor complex and associate with the co-SMAD (SMAD 4). The SMAD2/3/4 complex is translocated to the nucleus where, with cofactors, it alters expression of numerous genes including profibrotic genes such as collagens type I, III and V, plasminogen activator inhibitor −1 and connective tissue growth factor (Verrecchia et al., 2001; Chen et al., 2002). TGF-β receptor engagement can also result in activation of non-canonical signaling pathways including mitogen activated protein kinases, Rho-like GTPases and phosphatidylinositol-3-kinase (PI3K)/ Akt. The downstream biological outputs of TGF-β signaling are highly dependent on the pathological and physiological cellular context (Morikawa et al., 2016). The diverse effects of TGF-β and the pathophysiological context-dependent responses present hurdles in targeting TGF-β signaling therapeutically.

Emodin is an anthraquinone derivative that is found primarily in several Chinese herbs. Emodin appears to possess an array of biological activities including modulation of immune and inflammatory processes, protection against neurological damage, and suppression of tumor growth and metastasis (Cha et al., 2015; Fang et al., 2019). Studies have illustrated that emodin treatment attenuates fibrosis in animal models including bleomycin-induced fibrosis of the lung (Chen et al., 2009), carbon tetrachloride-induced liver steatosis and fibrosis (Dong et al., 2009) and pressure overload-induced myocardial fibrosis (Xiao et al., 2019). Despite the spectrum of beneficial effects attributed to emodin, the underlying cellular and molecular mechanisms of these effects are not clear. In order to gain a better appreciation for the potential therapeutic use of emodin in myocardial fibrosis, studies were performed with isolated cardiac fibroblasts to assess the effects of emodin on activation, function and signaling of these cells in response to TGF-β1.

## Materials and Methods

### Cell isolation and culture

Adult (8-10 weeks of age) male Sprague Dawley rats (Harlan Laboratories, www.envigo.com) were used for isolation of cardiac fibroblasts (Stewart et al., 2010). All experiments with animals were performed in accordance with institutional guidelines and were approved by the University of South Carolina Institutional Animal Care and Use Committee (IACUC). Following acclimation in the University of South Carolina School of Medicine animal facility for 1 to 2 days, rats were anesthetized by inhalation of isoflurane and euthanized by cervical dislocation. Hearts were removed, rinsed in sterile saline and cardiac tissue was minced into 2-3 mm^3^ pieces. Cardiac tissue was digested with Liberase (Roche Life Sciences, www.lifescience.roche.com) at 37°C. Cells were allowed to adhere to culture dishes and fibroblasts purified by differential adhesion. Cells were cultured in Dulbecco’s Modified Eagle’s Medium (DMEM) containing 10% fetal bovine serum (GE Healthcare Life Sciences, www.cytivalifesciences.com) and antibiotics (hereafter referred to as “normal fibroblast medium”). Fibroblasts were passaged at 70 to 80 percent confluence, following incubation in trypsin/ ethylenediaminetetraacetic acid solution. Only fibroblasts at low passage number (less than 5) were used in assays as fibroblast phenotype diverges at high passage numbers (Stewart et al., 2010). Prior to treatment, culture medium was changed to DMEM containing 1.5% fetal bovine serum (low serum medium) for 24 h to reduce spontaneous conversion to a myofibroblast phenotype in response to serum components.

### Collagen gel contraction

Following culture for 24 h in low serum medium, cells were trypsinized and resuspended in low serum medium. Three-dimensional collagen hydrogels were formed by the addition of fibroblasts to bovine collagen type I (PureCol; Advanced BioMatrix, Inc., www.advancedbiomatrix.com) at a ratio of 50,000 cells per milliliter of collagen (1.2 mg/ml collagen concentration). Following polymerization at 37°C for 1 h, the 3-dimensional collagen hydrogels containing fibroblasts were dislodged from the plastic wells. Culture was continued for 24 h in low serum medium with 0 or 5 ng/ml TGF-β1 (R&D Systems, www.rndsystems.com) with emodin (0, 10 or 20 μM). After 24 h, the perimeter of the collagen gels was measured. Data are presented as the degree of collagen hydrogel contraction relative to untreated controls.

### Immunoblotting

Fibroblasts were cultured for 24 h in low serum medium as indicated above and culture then continued an additional 24 h in low serum medium containing 0 or 5 ng/ml TGF-β1 and emodin (0, 10 or 20 μM). Following treatment, cells were extracted in RIPA solution (150 mM sodium chloride, 1% Triton X 100, 0.5% deoxycholate, 0.1% sodium dodecyl sulfate, 1.5 mM ethylenediaminetetraacetic acid, 50 mM Tris, pH 8.0) containing protease inhibitors (Protease Inhibitor Tablets; ThermoFisher Scientific, www.thermofisher.com). Fibroblast lysates were centrifuged to remove insoluble material and protein concentration of the supernatants determined with the bicinchronic acid (BCA) protein assay (ThermoFisher Scientific, www.thermofisher.com). Following separation by sodium dodecyl sulfate – polyacrylamide gel electrophoresis (SDS-PAGE), proteins were transferred to nitrocellulose. Nitrocellulose membranes were rinsed in tris-buffered saline containing TWEEN 20 (TBS-T) and nonspecific binding blocked by incubation in TBS-T containing 5% powdered milk. Nitrocellulose was incubated in primary antisera overnight at 4°C, rinsed in TBS-T and incubated in secondary antibodies conjugated to horseradish peroxidase. Primary antisera included anti-alpha 1 integrin (RRID:AB_91199), anti-alpha 2 integrin (RRID:AB_11212896) and anti-beta 1 integrin (RRID:AB_91150) all from Millipore (MilliporeSigma, St. Louis, MO, www.emdmillipore.com). Immunoblots were rinsed in TBS-T, developed with the Pierce SuperSignal western blot detection reagent (ThermoFisher Scientific, www.thermofisher.com) and exposed to x-ray film. Images of x-ray films were taken using a BioRad GelDoc system and quantified with Alpha Innotech AlphaView software. Quantitative data from proteins of interest were normalized to glyceraldehyde phosphate dehydrogenase (RRID:AB_641107, Santa Cruz Biotechnology, Dallas, TX www.scbt.com) and data presented as fold of vehicle-treated controls.

For analyses of collagen type I and collagen type III accumulation, fibroblasts were treated as above and conditioned medium collected after 24 h of treatment. Media were centrifuged to remove cellular debris and the protein concentration was determined with the bicinchronic acid (BCA) protein assay (ThermoFisher Scientific, www.thermofisher.com). Following separation of proteins by SDS-PAGE, western blots were carried out as described above with antibodies against collagen type I (RRID:AB_638601, Santa Cruz Biotechnology, Dallas, TX, www.scbt.com) and collagen type III (RRID:AB_2082354, Santa Cruz Biotechnology, Dallas, TX, www.scbt.com). Immunoblots were rinsed in TBS-T, developed with the Pierce SuperSignal western blot detection reagent (ThermoFisher Scientific, www.thermofisher.com) and exposed to x-ray film. Images of x-ray films were taken using a BioRad GelDoc system and quantified with Alpha Innotech AlphaView software (ProteinSimple, San Jose, CA www.proteinsimple.com). Since extracellular collagen was assessed in the present studies, quantification of collagen proteins was normalized to total protein via fast green-stained blots (Alsridge et al., 2008; Gomes et al., 2014) and data presented as fold of vehicle-treated controls.

Due to the rapidity of TGF-β1 signal transduction pathway activation, cells to be analyzed for signaling pathways were pretreated for 24 h with varying doses of emodin (0, 10 or 20 μM). Fibroblasts were subsequently treated with 0 or 5 ng/ml of TGF-β1 for 1 h in the continued presence of the respective emodin dose. Cells were gently detached from culture plates and collected by centrifugation. The cell pellets were rinsed in phosphate-buffered saline and dissolved in M-PER mammalian protein extraction reagent (ThermoFisher Scientific, www.thermofisher.com) containing mini protease inhibitor cocktail (Sigma Chemicals). The cell lysate was centrifuged to remove insoluble material, the supernatant collected and total protein concentration determined. Protein samples were resolved by SDS-PAGE, transferred electrophoretically to nitrocellulose membranes and subsequently probed with primary antisera to SMAD2, pSMAD2, SMAD3, pSMAD3, p38, pp38, ERK1/2 or pERK1/2 (all from Cell Signaling Technology Inc, Danvers, MA, www.cellsignal.com). Nitrocellulose membranes were rinsed and incubated in horseradish peroxidase- conjugated anti-rabbit or anti-mouse serum as appropriate. Blots were developed in Clarity Western ECL detection reagent (BioRad Laboratories, www.bio-rad.com) and exposed to X-OMAT AR films (Eastman Kodak, Rochester, NY, www.kodak.com). Duplicate immunoblots were probed with anti-β-actin serum (RRID: AB_476697, Sigma-Aldrich, St. Louis, MO, www.sigmaaldrich.com) as a loading control. Data are presented as the levels of the phosphorylated protein relative to its respective non-phosphorylated form.

### Migration

A wound healing assay (Liang et al., 2007) was utilized to assay the effects of TGF-β1 and emodin on heart fibroblast migration. Confluent cells grown in 6-well plates were cultured for 24 h in low serum medium. A scratch was made in the center of the confluent fibroblast culture with a 200-µl pipette tip. The cells were gently rinsed with sterile Moscona’s solution and culture continued in low serum medium containing 0 or 5ng/ml TGF-β1 and emodin (0, 10 or 20 μM). After 24 h of culture, images were captured on an inverted microscope and the relative migration of fibroblasts into the denuded area was measured using ImageJ (National Institutes of Health; Bethesda, MD).

### Proliferation

Proliferation of cardiac fibroblasts was assessed by a 5′-bromo-2′-deoxyuridine (BrdU) incorporation assay. Glass coverslips were coated with with 10 μg/ml collagen I to promote cell adhesion (Advanced BioMatrix, Inc, www.advancedbiomatrix.com). Following culture in low serum medium, cells were incubated for 24 h with 0 or 5ng/ml TGF-β1 and emodin (0, 10 or 20 μM). Culture medium contained BrdU (20 μM) during the treatment period. Coverslips were rinsed and fixed in absolute ethanol containing 10 mM glycine (pH 2.0) at −20°C for 30 m. BrdU incorporation was detected by immunocytochemical staining (Roche Applied Science, www.lifescience.roche.com). Following immunocytchemical staining with anti-BrdU, cells were co-stained with 4′-6-diamidino-2-phenylindole (DAPI) to visualize all cell nuclei. Cells were examined by using a Nikon E600 fluorescent microscope and the ratio of BrdU positive cells to total cell numbers were determined.

### Statistical Analysis

All experiments were repeated at least in triplicate with independent sets of fibroblasts as indicated in figure legends. Quantitative data were plotted in GraphPad Prism and statistical analyses carried out by one-way ANOVA followed by post-hoc Dunnett's test (multi-group comparison).

## Results

### Extracellular matrix expression

As mentioned above, emodin has been shown to reduce extracellular matrix accumulation in animal models of fibrosis. To evaluate this in heart fibroblasts, cells were treated for 24 h with TGF-β1 (0 or 5 ng/ml) and varying concentrations of emodin (0, 10 or 20 μM) followed by examination of collagen type I and collagen type III protein levels in the conditioned medium by immunoblot analysis. These experiments illustrated a significant increase in the levels of collagen type I (Figure 1A) and a slight but insignificant increase in collagen type III (Figure 1B) in conditioned medium of cardiac fibroblasts following treatment with TGF-β1 alone. The levels of both collagen types were reduced in a concentration-dependent manner by emodin relative to TGF-β1 treatment alone. Treatment with emodin also significantly reduced the basal levels (in the absence of TGF-β1) of collagen type I in the conditioned medium with a slight but statistically insignificant reduction in collagen type III.

**Fig. 1.**
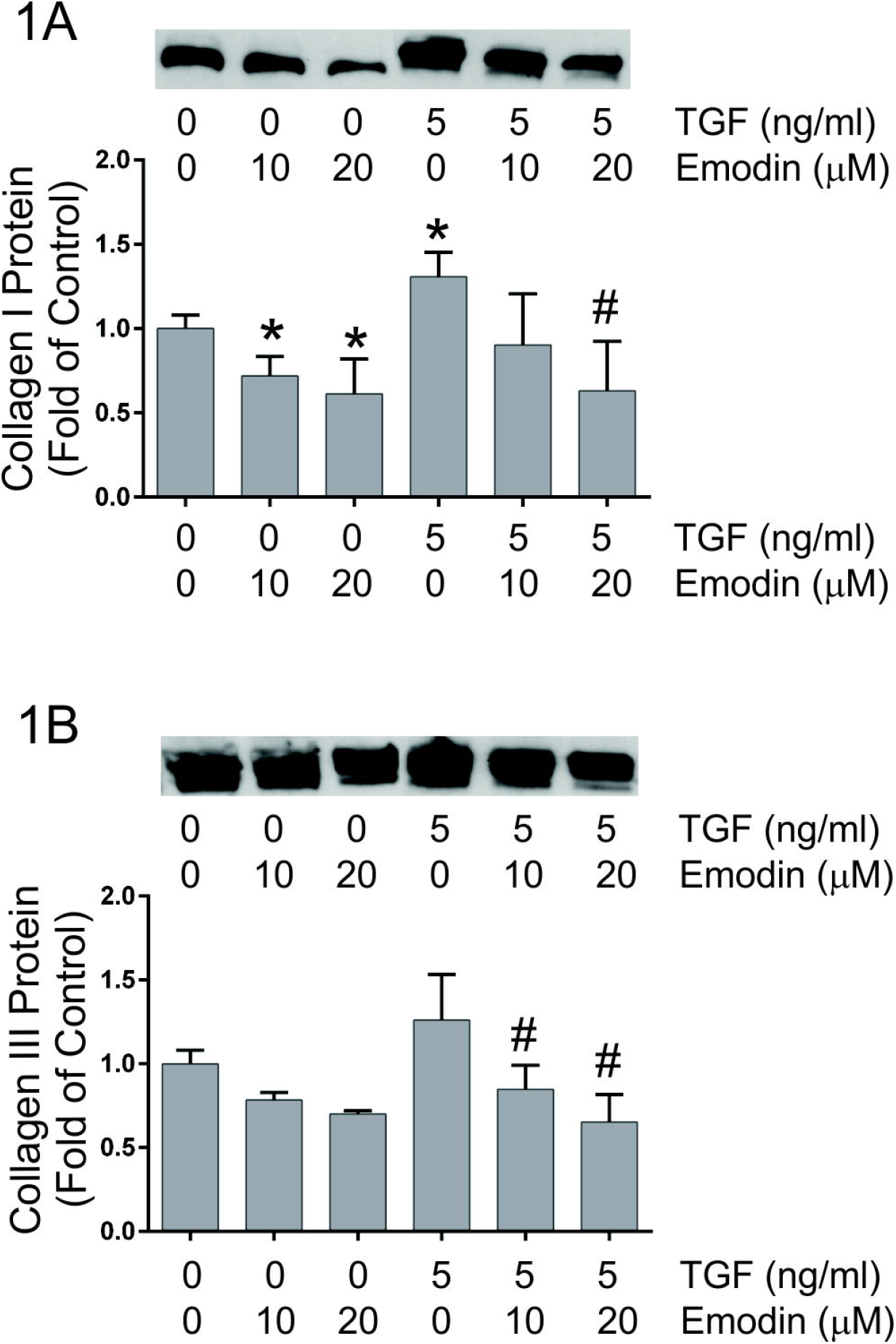
This illustrates relative quantification of the effects of TGF-β1 and emodin on collagen type I (Figure 1A) and collagen type III (Figure 1B) protein accumulation in conditioned medium of cardiac fibroblasts as determined by immunoblot analyses. The insets illustrate representative immunoblots with the anti-collagen type I and anti-collagen type III sera. (* p<0.05 relative to untreated controls, # p <0.05 relative to TGF-β1 treatment alone as determined by ANOVA, n = 4).

### Fibroblast bioassays

Exposure of several types of cells to TGF-β1 results in transformation or differentiation to a myofibroblast phenotype (Carthy et al., 2015; Pattarayan et al., 2018). Myofibroblasts have increased contractile activity, which we measured indirectly via their ability to contract 3-dimensional collagen hydrogels. Treatment of isolated adult heart fibroblasts with TGF-β1 resulted in enhanced contraction of three-dimensional collagen hydrogels (Figure 2). This response to TGF-β1 has been demonstrated previously (27, 68). Simultaneous treatment with emodin attenuated TGF-β1-induced collagen hydrogel contraction. The highest dose of emodin used in these studies had a slight but insignificant effect on basal collagen hydrogel contraction by cardiac fibroblasts (Figure 2).

**Fig. 2.**
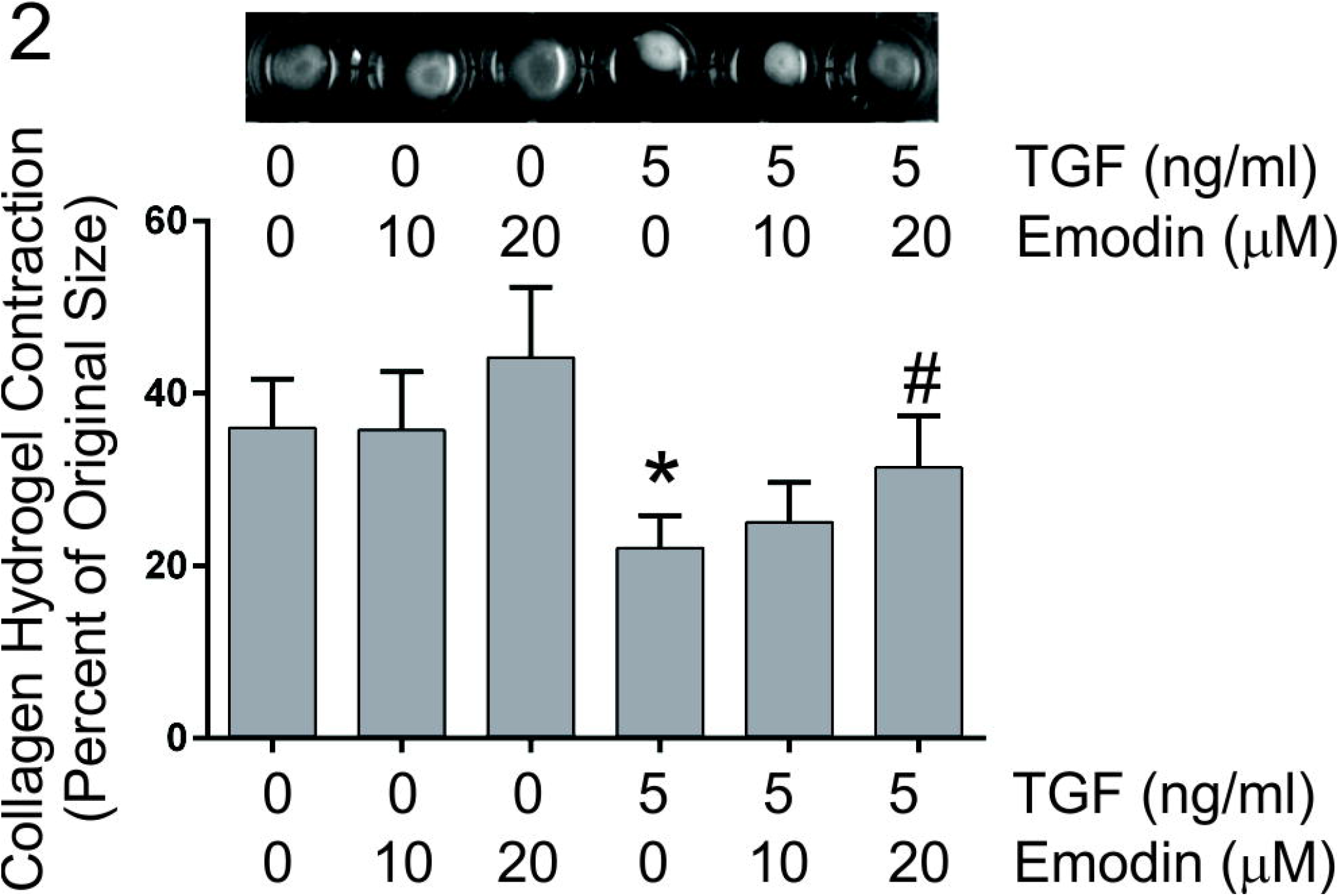
This figure demonstrates the effects of TGF-β1 and emodin on 3-dimensional collagen hydrogel contraction. Data are presented as the size of the hydrogel perimeter relative to the starting size (shorter bars indicate greater contraction). The inset shows representative images of collagen hydrogels. (* p<0.05 relative to untreated controls, # p <0.05 relative to TGF-β1 treatment alone as determined by ANOVA, n = 4).

Remodeling and contraction of collagen hydrogels can be impacted not only by the contractility of the cells but also by migration and proliferation of cells within the 3-dimensional collagen scaffold. Bioassays were performed to evaluate the effects of TGF-β1 and emodin on these aspects of fibroblast function. Scratch wound healing assays were carried out to assess the effects of TGF-β1 and emodin on fibroblast migration. Treatment of cells with 5ng/ml TGF-β1 had little effect on the ability of rat cardiac fibroblasts to repopulate a denuded area through a confluent culture (Figure 3A). Emodin treatment resulted in reduced ability of fibroblasts to repopulate the denuded area of the culture under basal conditions or when stimulated with TGF-β1 (Figure 3A). As repopulation of the wounded area of a confluent culture can occur through migration of the cells into the denuded area or proliferation of cells, the impact of TGF-β1 and emodin on fibroblast proliferation was assessed by BrdU incorporation. TGF-β1 treatment resulted in increased BrdU incorporation (Figure 3B) relative to untreated fibroblasts. In contrast to studies with an alveolar epithelial cell line (Gao et al., 2017), simultaneous treatment of cardiac fibroblasts with emodin had no detectable effect on basal proliferation nor the proliferative response to TGF-β1 (Figure 3B).

**Fig. 3.**
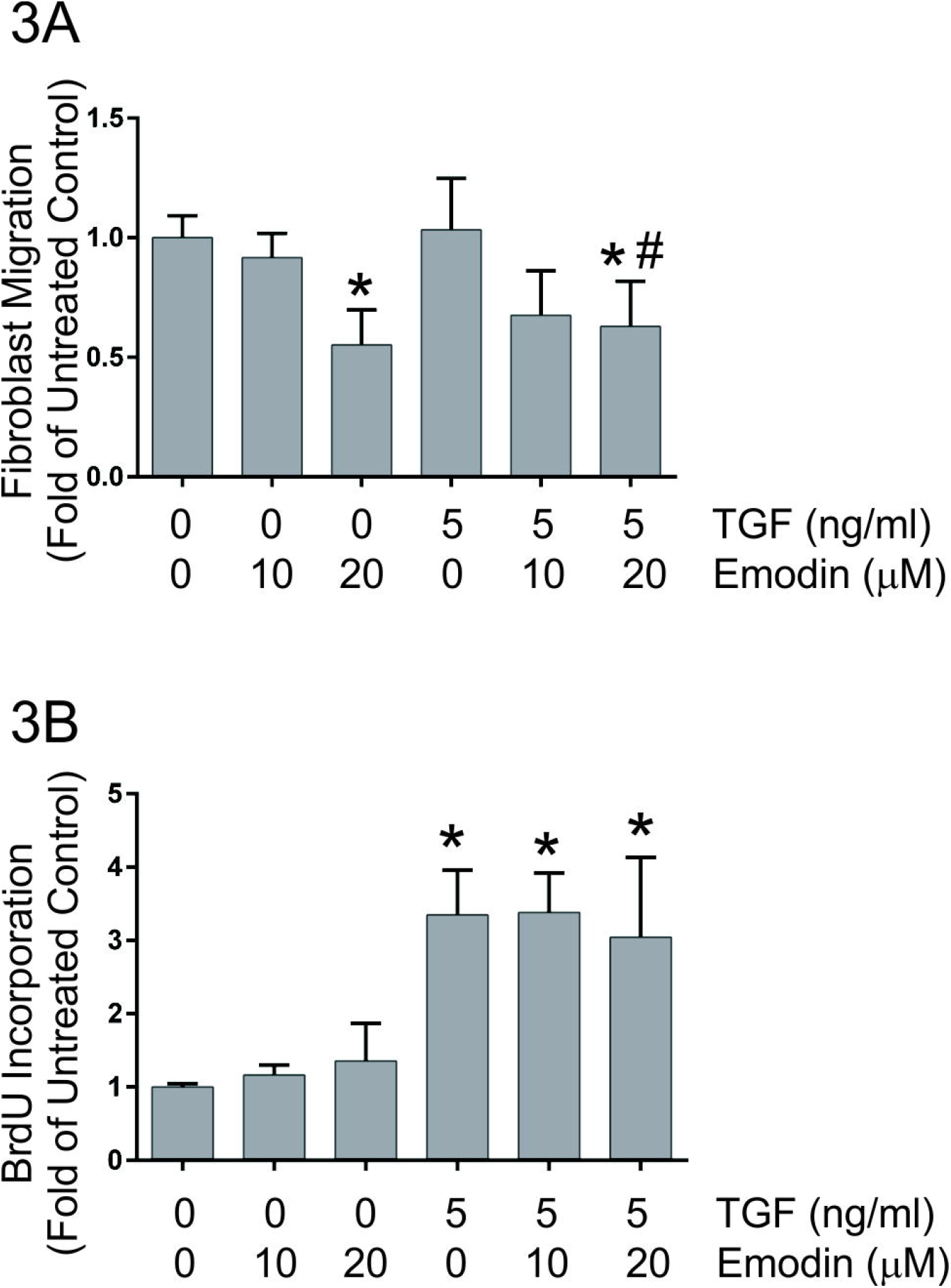
This demonstrates the effects of TGF-β and emodin on fibroblast migration (Figure 3A) and proliferation (Figure 3B). Data are presented as fold of the untreated controls. (* p <0.05 relative to untreated controls, # p <0.05 relative to TGF-β1 treatment alone as determined by ANOVA, n = 4).

### Fibroblast activation

The levels of α-smooth muscle actin and fibroblast activation protein, widely used as markers of myofibroblast phenotype, were evaluated by immunoblots following 24 h of treatment with 0 or 5 ng/ml TGF-β1 in the presence of 0, 10 or 20 μM emodin. Immunoblot analysis indicated increased levels of α-smooth muscle actin (Figures 4A and 4B) and fibroblast activation protein (Figures 4A and 4C) in fibroblast lysates following TGF-β1 treatment. Similar to recent studies demonstrating that emodin attenuates angiotensin II-induced fibroblast activation (Xiao et al., 2019), simultaneous treatment of fibroblasts with emodin and TGF-β1 reduced α-smooth muscle and fibroblast activation protein levels compared to treatment with TGF-β1 alone. Emodin treatment at the highest dose evaluated (20 μM) also significantly reduced α-smooth muscle actin protein levels under basal conditions.

**Fig. 4.**
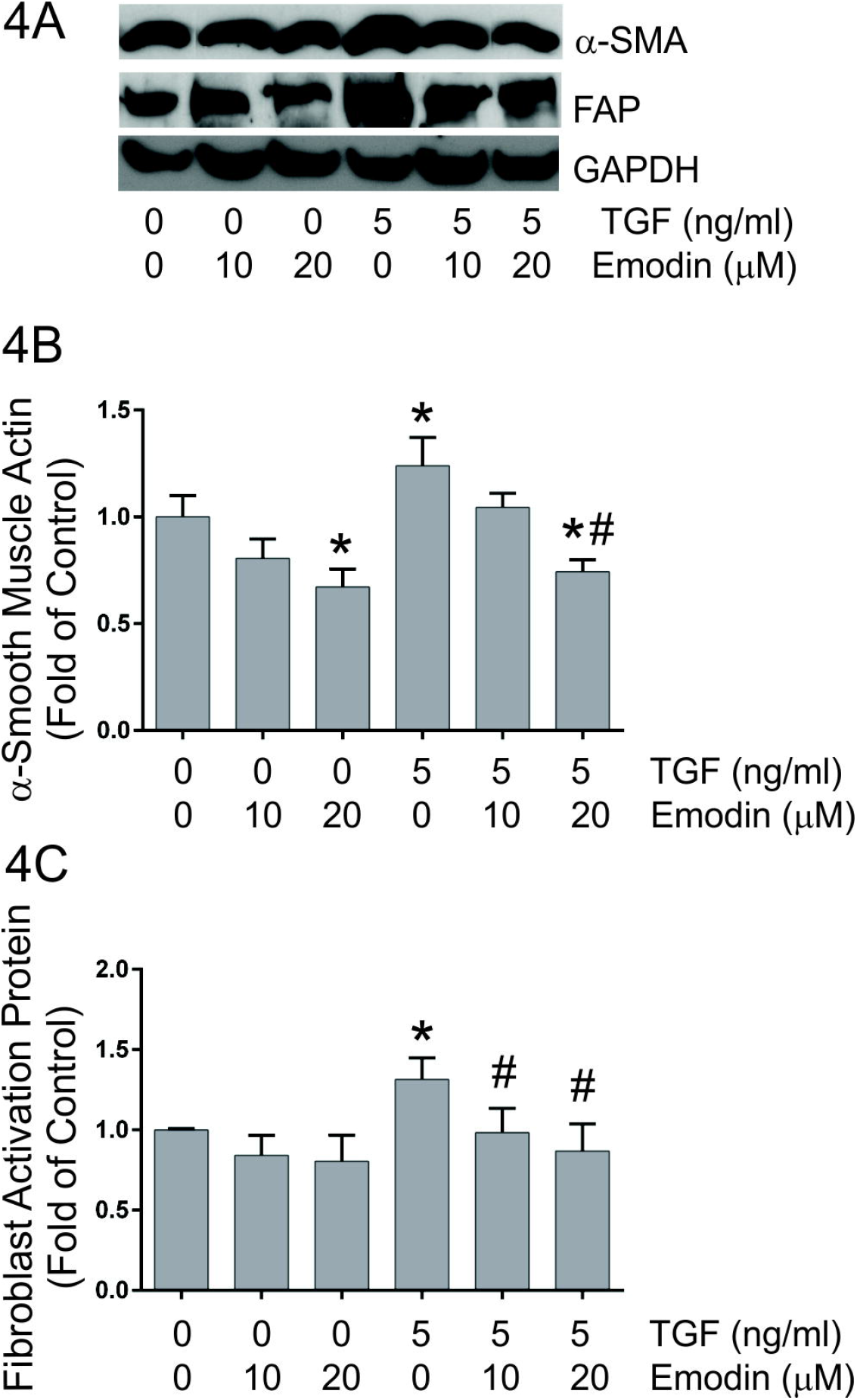
This figure illustrates the quantitative effects of TGF-β and emodin on the expression of α-smooth muscle actin (Figure 4B) and fibroblast activation protein (Figure 4C) levels as determined by immunoblots of cellular lysates. Data are presented as fold of the untreated controls. (* p <0.05 relative to untreated controls, # p <0.05 relative to TGF-β1 treatment alone as determined by ANOVA, n = 4). Images of representative immnoblots are shown in Figure 4A (α-SMA is α-smooth muscle actin, FAP is fibroblast activation protein and GAPDH is glyceraldehyde phosphate dehydrogenase).

### Integrin expression

As cell surface receptors for extracellular matrix components, integrins play pivotal roles in a number of cellular processes including remodeling and contraction of 3-dimensional collagen hydrogels (Gullberg et al., 1990; Carver et al., 1995). As previously illustrated (Fix et al., 2019), treatment of isolated adult rat heart fibroblasts with TGF-β1 resulted in significantly increased levels of α1 and α2 integrin proteins in the cells (Figures 5A - 5C, respectively). Treatment with TGF-β1 resulted in a slight but insignificant increase in β1 integrin protein in the cardiac fibroblasts (Figure 5D). Simultaneous treatment with emodin attenuated the TGF-β1-induced increases in α1 and α2 integrin protein levels and the highest dose of emodin reduced α1 integrin protein levels under basal conditions (Figures 5B and 5C).

**Fig. 5.**
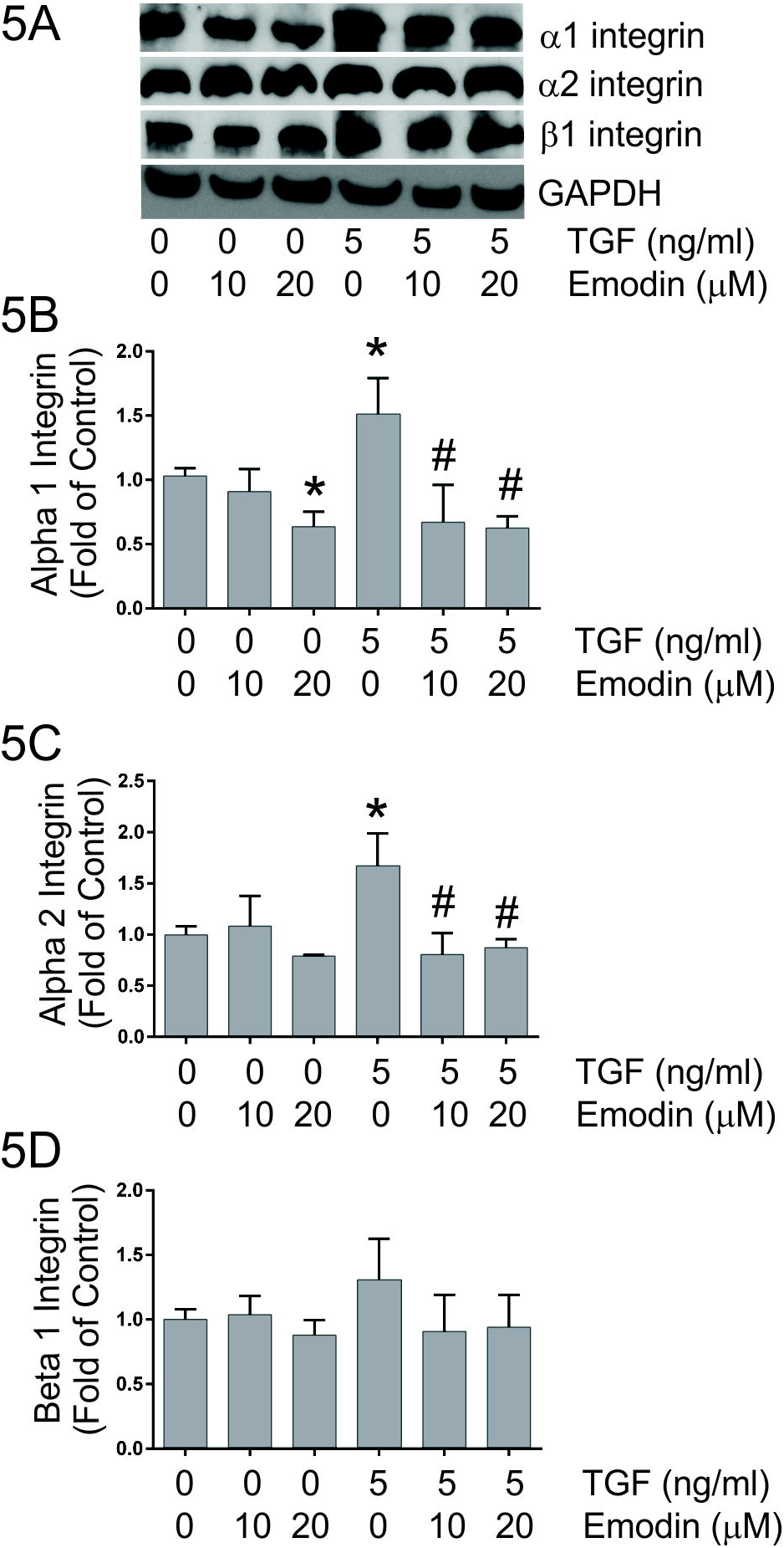
This figure demonstrates the effects of TGF-β1 and emodin on the levels of α1 integrin (Figure 5B), α2 integrin (Figure 5C) and β1 integrin (Figure 5D) protein levels as determined by immunoblots of cellular lysates. Data are presented as fold of the untreated controls. (* p <0.05 relative to untreated controls, # p <0.05 relative to TGF-β1 treatment alone as determined by ANOVA, n = 4). Images of representative immunoblots are shown in Figure 5A.

### Matrix metalloproteinase expression

Matrix metalloproteases (MMP) play critical roles in remodeling of the cardiac extracellular matrix (Lindsey, 2019). Conditioned medium was analyzed to evaluate the effects of TGF-β1 and emodin on the major MMPs produced by cardiac fibroblasts. Treatment of fibroblasts isolated from adult rat hearts with TGF-β1 had no apparent effect on MMP2 or MMP9 protein levels relative to untreated controls. Treatment of cardiac fibroblasts with emodin resulted in enhanced MMP2 and MMP9 protein levels, regardless of the presence or absence of TGF-β1 (Figures 6A and 6B).

**Fig. 6.**
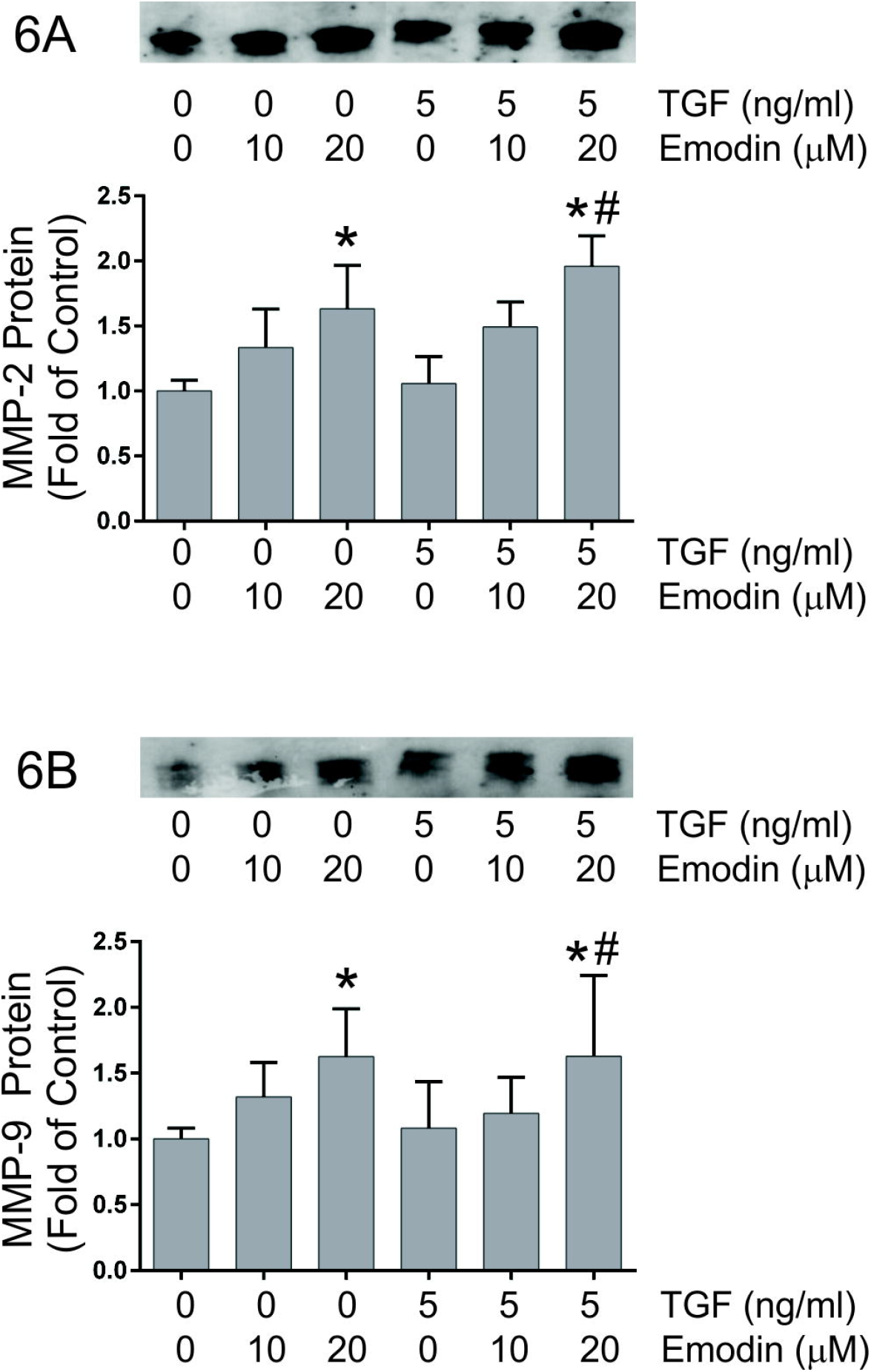
Graphic representation of the effects of TGF-β1 and emodin on the levels of matrix metalloprotease 2 (Figure 6A) and matrix metalloprotease 9 (Figure 6B) protein levels as determined by immunoblots of conditioned medium. The insets illustrate representative immunoblots with the respective matrix metalloprotease antisera. Data are presented as fold of the untreated controls. (* p <0.05 relative to untreated controls, # p <0.05 relative to TGF-β1 treatment alone as determined by ANOVA, n = 4).

### Activation of TGF-β-related signaling pathways

A number of signaling pathways are activated by exposure to TGF-β including the canonical SMAD2/3 pathway and non-canonical pathways. Immunoblots were utilized to evaluate the effects of emodin on activation of signaling pathways by TGF-β1 in heart fibroblasts. These studies demonstrated that treatment of fibroblasts with 5 ng/ml of TGF-β1 resulted in enhanced activation of the canonical TGF-β signaling pathway indicated by increased phosphorylation of SMAD2 (Figure 7A) and SMAD3 (Figure 7B). These were inhibited to approximately baseline levels by simultaneous treatment with emodin and TGF-β1. TGF-β1 treatment also activates non-canonical signaling pathways in cardiac fibroblasts indicated by significantly increased levels of phosphorylated forms of Erk 1/2 (Figure 7C); however, TGF-β1 treatment had no effect on p38 phosphorylation (Figure 7D). Treatment of cardiac fibroblasts with emodin attenuated the activation of Erk 1/2 by TGF-β1. Interestingly, simultaneous treatment of isolated heart fibroblasts with TGF-β1 and emodin resulted in a dose-dependent increase in p38 phosphorylation relative to TGF-β1 or emodin alone.

**Fig. 7.**
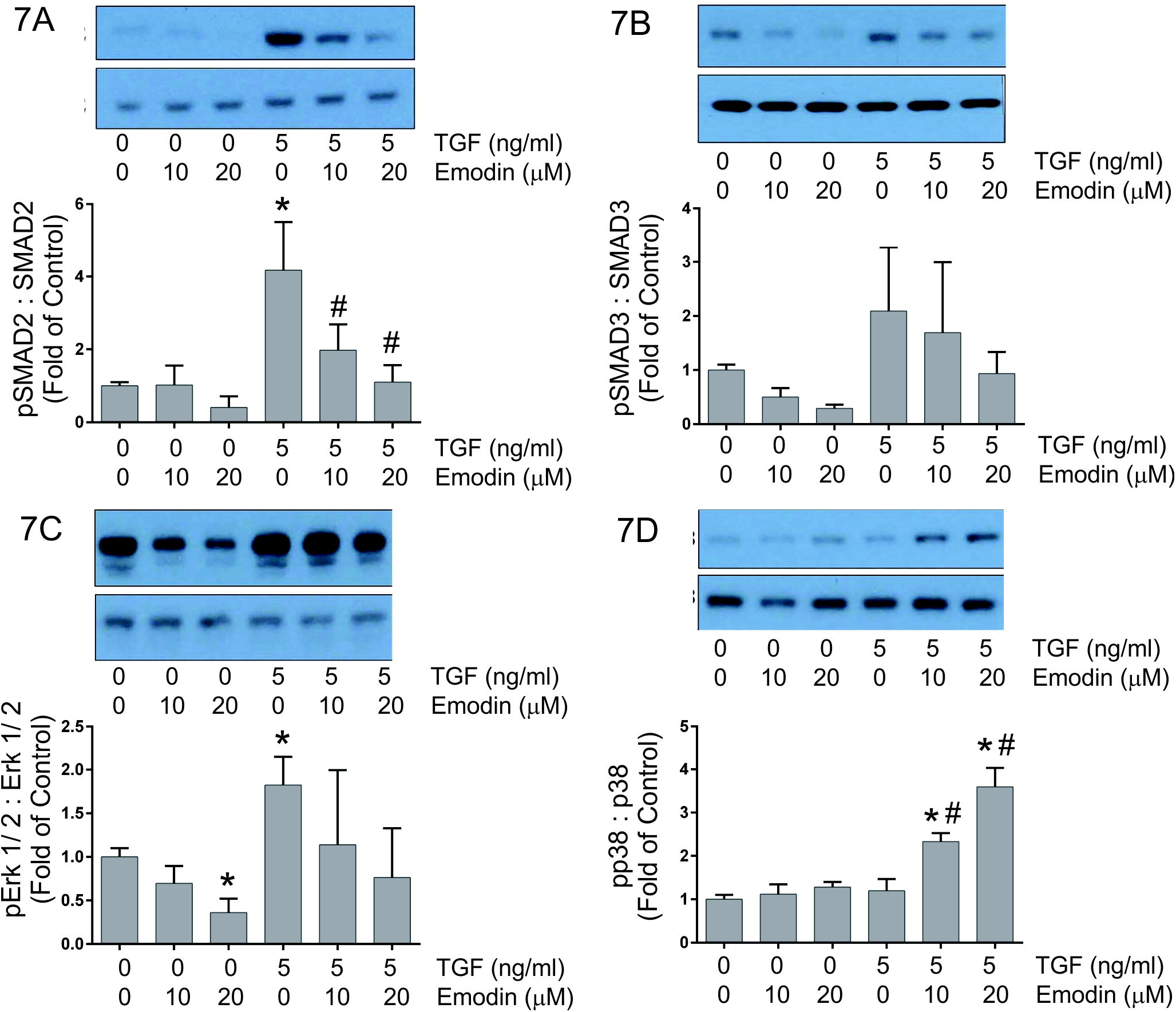
Graphic representation of the effects of TGF-β1 and emodin on the activation of TGF-β signaling components including SMAD 2 (Figure 7A), SMAD 3 (Figure 7B), Erk 1/ 2 (Figure 7C) and p38 (Figure 7D). The insets illustrate representative immunoblots with the respective signaling component antisera. Data are presented as the signal with antisera to the phosphorylated protein relative to antisera to total protein. (* p <0.05 relative to untreated controls, # p <0.05 relative to TGF-β1 treatment alone as determined by ANOVA, n = 3).

## Discussion

Fibrosis is an essential hallmark of adverse cardiac remodeling and is characterized by excessive deposition of extracellular matrix (ECM) and matricellular proteins (Spinale et al., 2016; Frangogiannis, 2019a). Cardiac fibrosis contributes to altered myocardial structure, geometry and compliance and has been implicated as an independent predictor of mortality in patients with non-ischemic heart failure (Aoki et al., 2011). Understanding the regulation of fibrosis has become increasingly important as it has been appreciated that in some settings the fibrotic response is reversible, at least in its early stages (Kumar and Sarin, 2007; Frangogiannis, 2019b). This realization has made the myofibroblast and other ECM-producing cells important therapeutic targets in fibrosis. Despite the substantial morbidity and mortality of fibrotic disease, only a handful of drugs have been approved by the FDA specifically for anti-fibrotic therapy. Recent reviews have highlighted some of the major hurdles to developing anti-fibrotic treatments which include the complexity of fibrotic signaling and undesirable side effects of therapeutics in non-diseased tissues. (Friedman et al., 2013). The goal of the present study was to evaluate the potential of emodin, a plant-derived anthraquinone, to attenuate TGF-β1-mediated cardiac fibroblast activation and fibrosis.

TGF-β and activation of its signaling pathways are leading contributors to fibrosis in multiple organs and in response to diverse stressors (Biernacka et al., 2011; Khalil et al., 2017). TGF-β1 is secreted by many cell types in a latent complex that can be activated by multiple mechanisms. Previous studies have illustrated that treatment with emodin reduces TGF-β1 expression in animal models of fibrosis (Chen et al., 2009; Dong et al., 2009; Ma et al., 2019). Emodin treatment *in vitro* has also been demonstrated to attenuate canonical TGF-β signaling and SMAD transcriptional activity (Tian et al., 2018; Wang et al., 2018). Similar to these latter studies, treatment of cardiac fibroblasts with emodin substantially reduced canonical TGF-β signaling illustrated by reduced SMAD 2 and SMAD 3 phosphorylation. The specific roles of attenuated canonical TGF-β signaling in the anti-fibrotic effects of emodin remain to be determined.

TGF-β receptor engagement can also result in activation of several non-canonical signaling pathways including phosphatidylinositol-3-kinase (PI3K)/ Akt, mitogen activated protein kinases (MAPK) and Rho-like GTPases (Zhang, 2009; Yeganeh et al., 2013). Recent studies have demonstrated that treatment of a hepatic stellate cell line with emodin *in vitro* reduces constitutive activation of p38 MAPK, while having no effect on ERK 1/ 2 or JNK (Wang et al., 2018). These effects on p38 activity were independent of SMAD signaling. Emodin treatment also attenuates p38, JNK and NF-κB activation following viral infection of MDCK cells with no effect on ERK 1/ 2 activity (19). In contrast to these results, treatment of cardiac fibroblasts with TGF-β1 resulted in ERK 1/2 activation, which was prevented by pretreatment with emodin. Interestingly, in the present study, emodin treatment, but only in the presence of TGF-β1, resulted in marked activation of p38. Future studies will be required to evaluate the functional effects of p38 activation by emodin in cardiac fibroblasts. Recent studies have shown that emodin can attenuate angiotensin II-induced fibroblast activation and that this is via upregulation of metastasis associated protein 3 (Xiao et al., 2019). It will be intriguing to determine whether modulation of metastasis associated protein 3 by emodin involves p38 activation.

There is debate regarding the effects of emodin on cell toxicity, proliferation and apoptosis. Several studies have illustrated that treatment of various cancer cell lines with emodin results in decreased proliferation (Cha et al., 2015; Sugiyama et al., 2019). This is in contrast to our results illustrating an increase in proliferation of fibroblasts isolated from adult rat hearts following treatment with TGF-β1; however, emodin had no effect on basal proliferation nor proliferation induced by TGF-β1 in these cells. A recent comparison of the effects of emodin on melanoma B16F10 and mouse embryonic fibroblasts suggests that emodin may have cell-specific effects as it inhibited proliferation of melanoma cells while having no effect on mouse embryonic fibroblast proliferation (Sugiyama et al., 2019). Along the same lines, treatment of human vascular smooth muscle cells with emodin resulted in significant inhibition of proliferation while emodin had little effect on proliferation of human vascular endothelial cells (Xu et al., 2018). Similar to the divergent effects on proliferation, differences exist in the effects of emodin on apoptosis. A number of studies have illustrated that emodin exposure induces apoptosis of HeLa, hepatocarcinoma, L02 liver cell line, fibroblasts and others (Ma et al., 2019; Xiong et al., 2019; Zheng et al., 2019). In contrast, emodin treatment has been demonstrated to attenuate hypoxia-induced apoptosis in H9C2 cardiomyocytes (Zhang et al., 2019) and apoptosis of podocytes induced by endoplasmic reticulum stress (Tian et al., 2018). These studies suggest that the effects of emodin on specific cell types need to be carefully evaluated and use of emodin therapeutically may need to be accompanied by cell-specific targeting mechanisms.

Emodin has been shown fairly consistently to inhibit migration and epithelial-to-mesenchymal transition and this, in part, forms the basis for its anti-cancer effects. Most of these studies have been performed with diverse cancer cell lines (Lin et al., 2016; Fang et al., 2019). Our studies herein with primary cardiac fibroblasts also demonstrate an inhibitory effect of emodin on migration using an *in vitro* wound healing assay. Diverse molecular mechanisms contributing to the inhibition of migration by emodin have been described suggesting the involvement of multiple pathways that may be cellular or context dependent. Several studies have demonstrated that emodin treatment diminishes VEGF levels or VEGF receptor signaling and that this functionally contributes to impaired migration (Dai et al., 2019). Other studies have illustrated that emodin treatment results in decreased expression of integrin linked kinase, an important player in migration and epithelial-to-mesenchymal transition (Lu et al., 2017). Our study demonstrates reduced levels of collagen-binding integrins (α1 and α2) in cardiac fibroblasts following emodin treatment. While functional studies were not performed, these integrins have been shown previously to be critical for fibroblast migration and extracellular matrix remodeling and are important intermediaries in cardiovascular disease (Carver et al., 1995; Chen et al., 2016).

## Acknowledgements

The authors would like to thank the University of South Carolina School of Medicine Instrumentation Resource Facility directed by Dr. Bob Price for their assistance in these studies.

## Data Availability Statement

All supporting data were illustrated and referred to in the manuscript as needed. The data that support the findings of this study are available from the corresponding author upon request.

## Literature Cited

Aldridge GM, Podrebarac DM, Greenough WT, Waller I (2008) J The use of total protein stains as loading controls: an alternative to high-abundance single protein controls in semi-quantitative immunoblotting. J Neurosci Meth 172:250–254. doi: 10.1016/j.jneumeth.2008.05.003.

Aoki T, Fukumoto Y, Sugimura K, Oikawa M, Satoh K, Nakano M, Nakayama M, Shimokawa H (2011) Prognostic impact of myocardial interstitial fibrosis in non-ischemic heart failure. Comparison between preserved and reduced ejection fraction heart failure. Circ J 75:2605–2613. doi: 10.1253/circj.cj-11-0568.

Biernacka A, Dobaczewski M, Frangogiannis NG (2011) TGF-β signaling in fibrosis. Growth Factors 29:196–202. doi: 10.3109/08977194.2011.595714.

Bhandary B, Meng Q, James J, Osinska H, Gulick J, Valiente-Alandi I, Sargent MA, Bhuiyan MS, Blaxall BC, Molkentin JD, Robbins J (2018) Cardiac fibrosis in proteotoxic cardiac disease is dependent upon myofibroblast TGF-β signaling. J Am Heart Assoc Oct 16;7(20):e010013. doi: 10.1161/JAHA.118.010013.

Branton MH, Kopp JB (1999) TGF-beta and fibrosis. Microbes Infect 1:1349–1365. doi: 10.1016/s1286-4579(99)00250-6.

Carthy JM (2018) TGFβ signaling and the control of myofibroblast differentiation: implications for chronic inflammatory disorders. J Cell Physiol 233:98–106. doi: 10.1002/jcp.25879.

Carthy JM, Sundqvist A, Heldin A, van Dam H, Kletsas D, Heldin CH, Moustakas A (2015) Tamoxifen inhibits TGF-β-mediated activation of myofibroblasts by blocking non-Smad signaling through ERK1/2. J Cell Physiol 230:3084–3092. doi: 10.1002/jcp.25049.

Carver W, Molano I, Reaves TA, Borg TK, Terracio L (1995) Role of the alpha 1 beta 1 integrin complex in collagen gel contraction in vitro by fibroblasts. J Cell Physiol 165:425–437. doi: 10.1002/jcp.1041650224.

Cha TL, Chuang MJ, Tang SH, Wu ST, Sun KH, Chen TT, Sun GH, Chang SY, Yu CP, Ho JY, Liu SY, Huang SM, Yu DS (2015) Emodin modulates epigenetic modifications and suppresses bladder carcinoma cell growth. Mol Carcinog 54:167–177. doi: 10.1002/mc.22084.

Chen C, Li R, Ross RS, Manso AM (2016) Integrins and integrin-related proteins in cardiac fibrosis. J Mol Cell Cardiol 93: 162–174. doi: 10.1016/j.yjmcc.2015.11.010.

Chen XH, Sun RS, Hu JM, Mo ZY, Yang ZF, Jin GY, Guan WD, Zhong NS (2009). Inhibitory effect of emodin on bleomycin-induced pulmonary fibrosis in mice. Clin Exp Pharmacol Physiol 36:146–153. doi: 10.1111/j.1440-1681.2008.05048.

Chen Y, Blom IE, Sa S, Goldschmeding R, Abraham DJ, Leask A (2002) CTGF expression in mesangial cells: involvement of SMADs, MAP kinase, and PKC. Kidney Int 62:1149–1159. doi: 10.1111/j.1523-1755.2002.kid567.x.

Dai G, Ding K, Cao Q, Xu T, He F, Liu S, Ju W (2019). Emodin suppresses growth and invasion of colorectal cancer cells by inhibiting VEGFR2. Eur J Pharmacol 15; 859:172525. doi: 10.1016/j.ejphar.2019.172525.

Derynck R, Budi EH (2019) Specificity, versatility and control of TGF-β family signaling. Sci Signal Feb 26; 12(570). doi:10.1126/scisignal.aav5183.

Dong MX, Jia Y, Zhang YB, Li CC, Geng YT, Zhou L, Li XY, Liu JC, Niu YC (2009) Emodin protects rat liver from CCl(4)-induced fibrogenesis via inhibition of hepatic stellate cell activation. World J Gastroenterol 15:4753–4762. doi: 10.348/wjg.15.4753.

Fang L, Zhao F, Iwanowycz S, Wang J, Yin S, Wang Y, Fan D (2019) Anticancer activity of emodin is associated with downregulation of CD155. Int Immunopharmacol 75:105763. doi: 10.1016/j.intimp.2019.105763.

Fix C, Carver-Molina A, Chakrabarti M, Azhar M, Carver W (2019) Effects of the isothiocyanate sulforaphane on TGF-β1-induced rat cardiac fibroblast activation and extracellular matrix interactions. J Cell Physiol 234:13931–13941. doi: 10.1002/jcp.28075.

Frangogiannis NG (2019) The extracellular matrix in ischemic and nonischemic heart failure. Circ Res 125:117–146. doi: 10.1161/CIRCRESAHA.119.311148.

Frangogiannis NG (2019) Can myocardial fibrosis be reversed? J Amer Coll Cardiol 73: 2283–2285. doi: 10.1016/j.jacc.2018.10.094.

Friedman SL, Sheppard D, Duffield JS, Violette S (2013) Therapy for fibrotic diseases: nearing the starting line. Sci Transl Med 5:167sr1. doi: 10.1126/scitranslmed.3004700.

Gao R, Chen R, Cao Y, Wang Y, Song K, Zhang Y, Yang J (2017) Emodin suppresses TGF-β1-induced epithelial-mesenchymal transition in alveolar epithelial cells through Notch signaling pathway. Toxicol Appl Pharmacol 318:1–7. doi: 10.1016/j.taap.2016.12.009.

Gomes R, Gilda JE, Gomes AV (2014) The necessity of and strategies for improving confidence in the accuracy of Western blots. Expert Reviews Proteomics 11:549–560. doi: 10.1586/14789450.2014.939635.

Gullberg D, Tingström A, Thuresson AC, Olsson L, Terracio L, Borg TK, Rubin K (1990) Beta 1 integrin-mediated collagen gel contraction is stimulated by PDGF. Exp Cell Res 186:264–272. doi: 10.1016/0014-4827(90)90305-t.

Hinz B (2016) The role of myofibroblasts in wound healing. Curr Res Transl Med 64:171–177. doi: 10.1016/j.retram.2016.09.003.

Kanisicak O, Khalil H, Ivey MJ, Karch J, Maliken BD, Correll RN, Brody MJ, J Lin SC, Aronow BJ, Tallquist MD, Molkentin JD (2016) Genetic lineage tracing defines myofibroblast origin and function in the injured heart. Nat Commun 7:12260. doi: 10.1038/ncomms12260.

Khalil H, Kanisicak O, Prasad V, Correll RN, Fu X, Schips T, Vagnozzi RJ, Liu R, Huynh T, Lee SJ, Karch J, Molkentin JD (2017) Fibroblast-specific TGF-β-Smad2/3 signaling underlies cardiac fibrosis. J Clin Invest 127:3770–3783. doi: 10.1172/JCI94753.

Kumar M, Sarin SK (2007) Is cirrhosis of the liver reversible. Indian J Pediatr 4:493–499. doi: 10.1007/s12098-007-0067-1.

Leask A, Abraham DJ (2004) TGF-β signaling and the fibrotic response. FASEB J 18:816–827. doi: 10.1096/fj.03-1273rev.

Liang CC, Park AY, Guan JL (2007) In vitro scratch assay: a convenient and inexpensive method for analysis of cell migration in vitro. Nat Protoc 2:329–333. doi: 10.1038/nprot.2007.30.

Lin W, Zhong M, Liang S, Chen Y, Liu D, Yin Z, Cao Q, Wang C, Ling C (2016) Emodin inhibits migration and invasion of MHCC-97H human hepatocellular carcinoma cells. Exp Ther Med. 12:3369–3374. doi: 10.3892/etm.2016.3793.

Lindsey ML (2018) Assigning matrix metalloproteinase roles in ischaemic cardiac remodeling. Nat Rev Cardiol 15: 471–479. doi: 10.1038/s41569-018-0022-z.

Lu J, Xu Y, Zhao Z, Ke X, Wei X, Kang J, Zong X, Mao H, Liu P (2017). Emodin suppresses proliferation, migration and invasion in ovarian cancer cells by down regulating ILK in vitro and in vivo. Onco Targets Ther 10:3579–3589. doi: 10.2147/OTT.S138217.

Ma C, Wen B, Zhang Q, Shao PP, Gu W, Qu K, Shi Y, Wang B (2019) Emodin induces apoptosis and autophagy of fibroblasts obtained from patient with ankylosing spondylitis. Drug Des Devel Ther 13:601–609. doi: 10.2147/DDDT.S182087.

Morikawa M, Derynck R, Miyazono K (2016) TGF-β and the TGF-β family: Context-dependent roles in cell and tissue physiology. Cold Spring Harb Perspect Biol May 2;8(5). doi: 10.1101/cshperspect.a021873.

Pakshir P, Hinz B (2018) The big five in fibrosis: Macrophages, myofibroblasts, matrix, mechanics and miscommunication. Matrix Biol 69:81–93. doi: 10.1016/j.matbio.2018.01.019.

Pattarayan D, Sivanantham A, Krishnaswami V, Loganathan L, Palanichamy R, Natesan S, Muthusamy K, Rajasekaran S (2018) Tannic acid attenuates TGF-β1-induced epithelial-to-mesenchymal transition by effectively intervening TGF-β signaling in lung epithelial cells. J Cell Physiol 233:2513–2525. doi: 10.1002/jcp.26127.

Shu DY, Lovicu FJ (2017) Myofibroblast transdifferentiation: The dark force in ocular wound healing and fibrosis. Prog Retin Eye Res 60:44–65. doi: 10.1016/j.preteyeres.2017.08.001.

Spinale FG, Frangogiannis NG, Hinz B, Holmes JW, Kassiri Z, Lindsey ML (2016) Crossing into the next frontier of cardiac extracellular matrix research. Circ Res 119:1040–1045. doi: 10.1161/CIRCRESAHA.116.309916.

Stewart JA Jr, Massey EP, Fix C, Zhu J, Goldsmith EC, Carver W (2010) Temporal alterations in cardiac fibroblast function following induction of pressure overload. Cell Tissue Res 340:117–126. doi: 10.1007/s00441-010-0943-2.

Sugiyama Y, Shudo T, Hosokawa S, Watanabe A, Nakano M, Kakizuka A (2019) Emodin as a mitochondrial uncoupler, induces strong decreases in adenosine triphosphate (APT) levels and proliferation of B16F10 cells, owing to their glycolytic reserve. Genes Cells 24:569–584. doi: 10.1111/gtc.12712.

Tian N, Gao Y, Wang X, Wu X, Zou D, Zhu Z, Han Z, Wang T, Shi Y (2018) Emodin mitigates podocytes apoptosis induced by endoplasmic reticulum stress through the inhibition of the PERK pathway in diabetic nephropathy. Drug Des Devel Ther 12:2195–2211. doi: 10.2147/DDDT.S167405.

Travers JG, Kamal FA, Robbins J, Yutzey KE, Blaxall BC (2016) Cardiac fibrosis: the fibroblast awakens. Circ Res 118:1021–1040. doi: 10.1161/CIRCRESAHA.115.306565.

Verrecchia F, Tacheau C, Schorpp-Kistner M, Angel P, Mauviel A (2001) Induction of the AP-1 members c-Jun and JunB by TGF-beta/Smad suppresses early Smad-driven gene activation. Oncogene 20:2205–2211. doi: 10.1038/sj.onc.1204347.

Wang X, Niu C, Zhang X, Dong M (2018) Emodin suppresses activation of hepatic stellate cells through p38 mitogen-activated protein kinase and Smad signaling pathways in vitro. Phytother Res 32:2436–2446. doi: 10.1002/ptr.6182.

Xiao D, Zhang Y, Wang R, Fu Y, Zhou T, Diao H, Wang Z, Lin Y, Li Z, Wen L, Kang X, Kopylov P, Shchekochikhin D, Zhang Y, Yang B (2019) Emodin alleviates cardiac fibrosis by suppressing activation of cardiac fibroblasts via upregulating metastasis associated protein 3. Acta Pharm Sin B 9:724–733. doi: 10.1016/j.apsb.2019.04.003.

Xiong G, Chen H, Wan Q, Dai J, Sun Y, Wang J, Li X (2019) Emodin promotes fibroblast apoptosis and prevents epidural fibrosis through PERK pathway in rats. J Ortho Surg Res 14: 319. doi: 10.1186/s13018-019-1357-9.

Xu K, Al-ani MK, Wang C, Qiu X, Chi Q, Zhu P, Dong N (2018) Emodin as a selective inhibitor of vascular smooth muscle cells versus endothelial cells suppress arterial intima formation. Life Sci 207:9–14. doi: 10.1016/j.lfs.2018.05.042.

Yeganeh B, Mukherjee S, Moir LM, Kumawat K, Kashani HH, Bagchi RA, Baarsma HA, Gosens R, Ghavami S (2013) Novel non-canonical TGF-β signaling networks: emerging roles in airway smooth muscle phenotype and function. Pulm Pharmacol Ther 26:50–63. doi: 10.1016/j.pupt.2012.07.006.

Zhang YE (2009). Non-Smad pathways in TGF-beta signaling. Cell Res 19: 128–139. doi: 10.1038/cr.2008.328.

Zhang X, Qin Q, Dai H, Cai S, Zhou C, Guan J (2019) Emodin protects H9c2 cells from hypoxia-induced injury by up-regulating miR-138 expression. Braz J Med Biol Res 52: e7994. doi: 10.1590/1414-431X20187994.

Zheng X-Y, Yang S-M, Zhang R, Wang S-M. Li G-B, Zhou S-W (2019) Emodin-induced autophagy against cell apoptosis through the PI3K/AKT/mTOR pathway in human hepatocytes. Drug Des Devel Ther 13:3171–3180. doi: 10.2147/DDDT.S204958.

